# Evidence of Filopodial translocation of Blastema associated microRNA rich Exosome like Extracellular Vesicles

**DOI:** 10.64898/2026.06.15.732514

**Authors:** Poovizhi Shanmugam, Murali Mohan Mishra, Samta Gupta, Manyata Makkar, Debasish Durgamadhab Mishra

## Abstract

Zebrafish (*Danio rerio*) possess remarkable regenerative capacity, making them an ideal model for studying the molecular mechanisms underlying tissue regeneration. In this article we report the identification of blastema linked exosome like extracellular vesicles (EVs) in zebrafish, that to the vesicles were plausibly being translocated in the proximo-distal axis through filipodia. We further thoroughly examined the exosome like EVs isolated from regenerating tissues of zebrafish caudal fins to characterize their nucleic acid cargo and evaluate their potential regulatory functions in regeneration. Caudal fins were amputated and allowed to regenerate and exosome like EVs isolated from blastema tissues displayed increased abundance compared to non-amputated controls. RNA sequencing identified a dynamic cluster of EV linked microRNAs (miRs). These differentially expressed miRs, including dre-miR-21, dre-miR-200b, dre-miR-218a and dre-let-7e were upregulated and associated with promoting proliferation, migration, differentiation, and tumour suppression pathways. Moreover, dre-miR-100, dre-miR-146a and dre-miR-200c regulated osteogenic differentiation, inflammatory signalling, epithelial-mesenchymal transition, and cell adhesion. Regeneration is generally believed to be coordinated only by local morphogen diffusion. Through this study it is indicative that filipodia bound EVs might have a pivotal role in long-range communication between blastema and the proximal tissues during the regeneration process. A detailed analyses of the miR targets and their validation could potentially lead to novel advancement and solutions in the field of regeneration and regenerative medicine in the near future.

## INTRODUCTION

Epimorphic regeneration is a remarkable biological process that can restore complex tissues following injury. In humans and other mammals, regenerative capacity is limited and tissue repair typically occurs primarily through fibrotic scar formation rather than complete regeneration, leading to functional deficits. In contrast, few lower vertebrates have the ability to restore complex structures and organs including limbs, fins and internal organs (Tanaka and Reddien, 2011). Understanding the cellular and molecular mechanisms underlying such regenerative responses will have implications for regenerative biology and therapeutic development.

Among these models, the zebrafish (Danio rerio), is an established model for studying regeneration, due to its accessibility and rapid regenerative capacity. Following injury or amputation, regeneration proceeds through well characterized stages including wound healing, the formation of blastema, proliferative outgrowth and tissue patterning. The blastema is a population of a stem cell-like progenitor cells which are arise from the dedifferentiation of cells at the wound bed. These blastema cells proliferate and re-differentiate to restore the tissue architecture and function (Seifert and Muneoka, 2018). Successful regeneration requires coordinated communication between blastema cells and surrounding tissues through an intracellular network of molecular pathways through paracrine exchange of cytokines, and growth factors (Sun and Irvine, 2014). The research findings shows that pathways such as canonical Wnt, BMP, FGF and Sonic hedgehog signalling play central roles in blastema formation and regeneration outgrowth in multiple vertebrate systems, including zebrafish (Stoick-Cooper, Moon and Weidinger, 2007). However, the mechanisms through which these regenerative signals are coordinated and distributed across regenerating tissues remain incompletely understood.

Extracellular vesicles (exosome like EVs) have emerged as important mediators of intercellular communication. Exosome like EVs are nanosized membrane-bound vesicles released by cells that carry diverse bioactive cargo, including proteins, lipids, RNAs, and microRNAs (miRNAs), that reflect the physiological state of their cells of origin (Sun and Chang, 2024). These vesicles are transported through biological fluids and deliver their cargo to recipient cells, thereby mediating paracrine signalling (Février and Raposo, 2004; Mead and Tomarev, no date). Exosome like EVs are commonly characterized by markers including tetraspanins (CD9, CD63, CD81), heat shock proteins (Hsp70 and Hsp90), and endosomal proteins such as Alix and TSG101 (Sabin and Kikyo, 2014). Increasing evidence suggests that EV-mediated signalling contributes to diverse physiological and pathological processes.

In regenerative biology, exosome like EVs derived from stem and progenitor cells have been implicated in tissue repair, wound healing, immune modulation and angiogenesis (Uminska *et al*., 2025; Zhou *et al*., 2025). Conversely, tumour-derived exosome like EVs contribute to tumour progression, metastasis, immune suppression, and therapy resistance (Bao *et al*., 2018; Ma *et al*., 2021; Ten *et al*., 2024). The growing recognition of EV function in tissue repair has not been extended to explore their role in epimorphic regeneration. In 2020, Ohgo et al. reported a notable study on demonstrating the presence of Exosome like EVs in regenerating zebrafish caudal fins using in vivo electroporation of exosomal markers (Ohgo *et al*., 2020). However, the molecular composition, transport mechanisms, and functional significance of exosome like EVs during regeneration are largely unknown.

Recent studies in developmental systems have highlighted the role of specialized membrane protrusions, including filopodia, cytonemes, and tunnelling nanotubes, in accelerating long-range intercellular communication (Sagar *et al*., 2015). These structures allows the directed transport of signalling molecules and in certain cases, vesicular cargo between cells (Sanders, Llagostera and Barna, 2013). Increasingly, such mechanisms are recognized as important mediators of tissue patterning and morphogen distribution during development. Whether similar structures contribute to EV-mediated communication during vertebrate tissue regeneration remains unclear.

In addition, exosome like EVs act as carriers of regulatory nucleic acids, particularly microRNAs (miRNAs), which are small non-coding RNAs controlling gene expression at the post-transcriptional level. miRNAs have been emerged as important regulators of regenerative processes (Ramachandran, Fausett and Goldman, 2010; Yin *et al*., 2012). Thatcher et al in 2008 demonstrated differential expression of several miRNAs during zebrafish caudal fin regeneration, including upregulation of miR-200b and downregulation of miR-203, miR-2, miR-338, and miR-301. These miRNAs regulate genes associated with pathways such as Wnt and BMP signalling (Thatcher *et al*., 2008). Such findings underscore the significance of miRNAs in regeneration. However, whether regenerative miRNAs observed in intercellular signalling communicate through EV-mediated transport remains uncleared.

Despite the emerging interest in understanding exosome like EVs in regeneration, the major challenge is the isolation of exosome like EVs from solid tissues. Unlike exosome like EVs derived from cultured cells or biofluids, tissue derived-exosome like EVs require mechanical or enzymatic tissue dissociation, which may introduce the risk of contamination from cellular debris, thus compromising vesicle integrity and variability in yield or purity. Therefore, the establishment of reliable methods for isolating exosome like EVs from regenerating tissues is essential for understanding their biological functions.

In the present study, we examined the presence, distribution, and molecular cargo of exosome like EVs during zebrafish caudal fin regeneration. To overcome the technical challenges associated with tissue-derived EV isolation, two isolation approaches such as centrifugal filter-based ultrafiltration and conventional ultracentrifugation were comparatively evaluated for particle recovery and vesicle integrity. Subsequently, exosome like EVs isolated from regenerating tissues were characterized and subjected to RNA extraction and sequencing to identify regeneration-associated EV cargo, particularly miRNAs. In addition, in situ visualization was performed to examine the spatial organization of exosome like EVs within regenerating tissues and to explore the possible involvement of filopodia-like structures in EV-mediated intercellular communication. Collectively, this study aims to provide insights into the potential role of exosome like EVs as regulators of coordinated signalling during epimorphic regeneration.

## Results

### Exosomes located in regenerating zebrafish fin

Amputated zebrafish caudal fins showed a regeneration of around 31.5 ± 3.2 % (mean ± SD) by the 10 days post amputation(dpa) (see Figure S1). Later, visualization of regenerating fins at 10dpa for tetraspanin-9-positive like EVs (green) using whole mount immunostaining revealed that EVs were highly localized in the regenerating fins compared with freshly amputated control fins (0 dpa). In control fins, EVs were observed as sparse and isolated solo granules or small island of EV cluster (») mainly around the amputated tip region (Figure. 1A). In contrast, regenerating fins exhibited increased EV abundance, forming dense EV conglomerates (Figure 1B-D). Moreover, EV-rich islands were associated with the cytoplasm of blastemal cells (*) and with newly laid extracellular matrix, aligning with the collagen fibre like structures (*§*). There is another more pertinent possibility. The EVs are being transported from the blastema in the proximal direction via cytonemal or tunnelling nanotube like filopodial cytoplasmic extensions. Literature, showed that one cellular cluster often reach out to other clusters using similar filopodial extensions for exchange of morphogens and sometimes EVs in invertebrate and vertebrate embryos.

**Figure 1.**
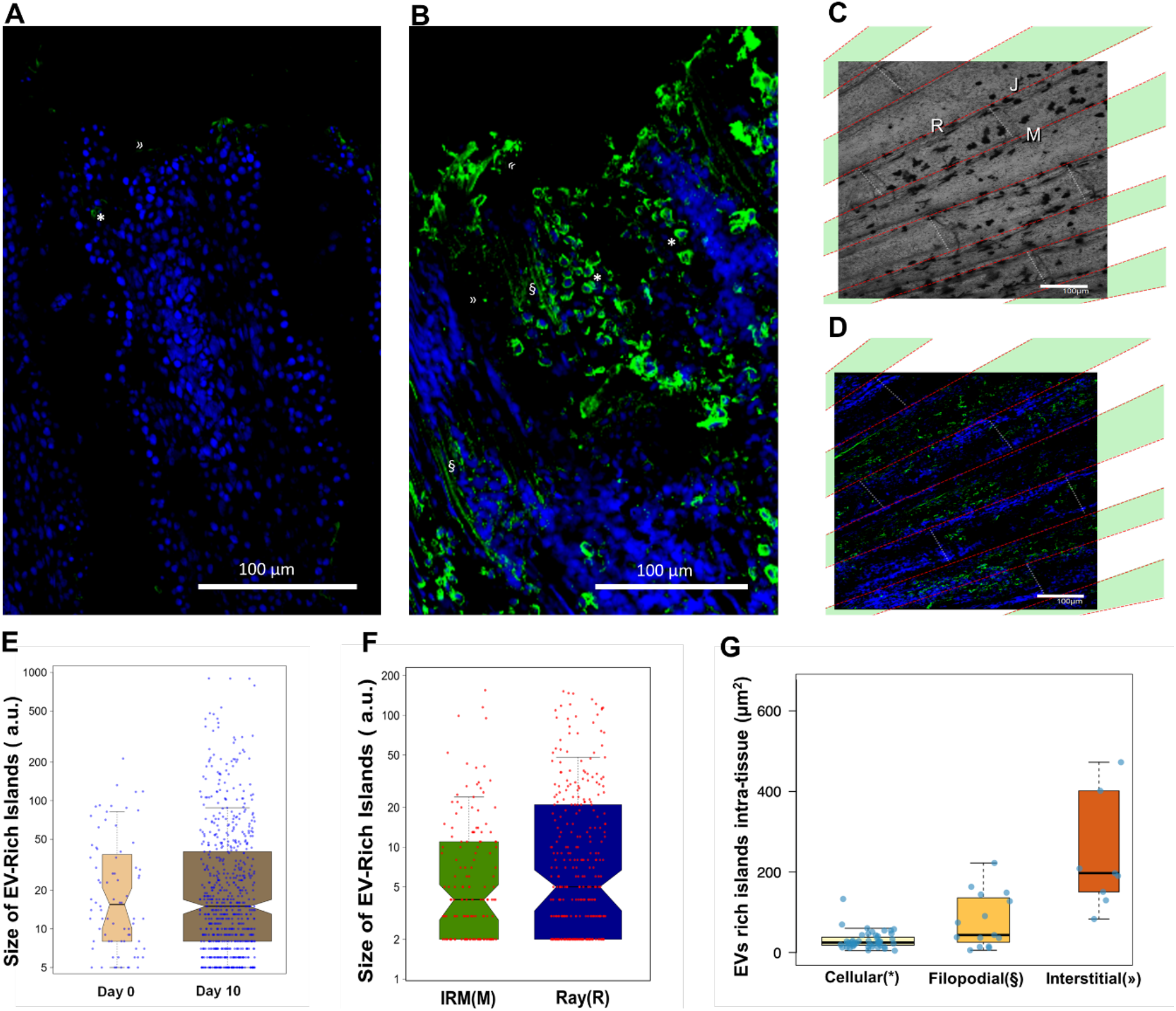
Whole mount confocal laser scanning microscopy of zebrafish caudal fin. EVs were stained with Alexa fluor 488 tagged anti-tetraspanin9 (or anti-CD63) antibody-green colour, and nuclei were counter stained with DAPI-blue colour. A- freshly amputated control (0 dpa), fin has not started regeneration; B-10 dpa regenerated caudal fin; C- DIC image of mid zone of regenerated fin; D- fluorescent image of mid zone of regenerated fin. E-The box plot with jittered data points of EV conglomerations or islands in the whole mount samples of 0 dpa and 10 dpa fins. The waistline of the notched box represents the median of the distribution. The width of the box is proportional to the number of data points, which is represented by blue coloured dots. Summation of the areas of all the EV-rich islands gives an estimate of total EV content. F- Comparison of total area of EV-rich islands in various zones of 10 dpa fins showed that the fin-rays had ∼two time more area than that of inter-ray membrane (IRM or M) zones. G- Comparison of total area of EV rich islands associated with cellular, filopodial and interstitial regions. *Note: (») solo granules or island of EV clusters, (*) cytoplasm associated EVs, and (§) points to filopodia associated EVs, R- caudal fin ray, J- bone joint of caudal fin rays, and M-inter-ray membrane*.

Such filopodial extension are special, and often characterized by their linearity and punctate appearance (Rustom *et al*., 2004; Sanders, Llagostera and Barna, 2013; Gradilla *et al*., 2014; Sagar *et al*., 2015; Daly, Hall and Ogden, 2022). In earlier reports, the puncta were shown to travel maintaining sufficient gaps between each other along the filopodial track. The EV containing puncta could be clearly differentiated from the filopodial track using GFP tagged CD63. However, in our case the gap between the puncta in the filopodial tracks seemed to be very short, as if they are tandem ‘railway-coaches’. If this is the case, it might be the first report that shows filopodia liked EV transport in regenerating tissues. However, the claim needs to be validated further before general acceptance. The colocalization of shh protein with the EVs in the puncta could also be demonstrated for further confirmation as shown earlier in reports (Gradilla *et al*., 2014).

Differential interference contrast (DIC) and fluorescence imaging of the mid-zone of regenerated fins shows that EV rich islands were found mostly around the regenerating caudal fin rays (R) and joints (J), whereas relatively fewer EVs were observed in inter ray membrane (M) (Figure.1C, D). Quantitative image analysis (Figure 1E) revealed that the total EV area increased from 2378 units in control fins to 29752 units in regenerating fins, indicating more than a 10-fold elevation of EV biogenesis during regeneration. Further analyses of EV distribution in regenerating fins (Figure 1F) revealed that EV-rich islands were increased two-fold in fin rays compared with inter-ray membrane zones. This observation was indicative of a substantial exchange of EV between distal blastema and proximal portion of the regenerating caudal fin, particularly concentrated in bone associated regions (fin rays). To further characterize the spatial organization of EVs, EV positive regions were classified into cell associated, filopodial and interstitial EV populations. Quantitative analysis demonstrated distinct differences in EV-positive area among these categories (Figure 1G). Cell-associated EVs were predominantly observed as small punctate structures, whereas filopodial EVs exhibited intermediate-sized regions with a broader range of area values. Interstitial (free) EV-rich islands occupied the largest EV-positive areas, with some regions exceeding 400 µm^2^. Collectively, these observations indicate an increasing trend in EV-positive area from cellular to filopodial and interstitial compartments. Together, these clarifies that epimorphic regeneration is not superficial. However, the directionality of this exchange was not resolved from the data and required further investigation.

### Regenerating fin exosomes have typical shape and size

Once densely packed EVs were observed in regenerating fins, attempts were made to isolate and characterize them. Low-cost ultrafiltration (UF) and ultracentrifugation (UC) methods were employed for isolation of EVs. The isolated EVs were characterized for their morphology and structural integrity using field emission scanning electron microscope (FESEM) and transmission electron microscope (TEM) respectively (Figure. 2). In both methods, vesicle size ranged around 200 nm which was consistent with the characteristic dimension of exosomes given by MISEV guidelines (Théry *et al*., 2018; Welsh *et al*., 2024) (see Figure S2).

**Figure 2.**
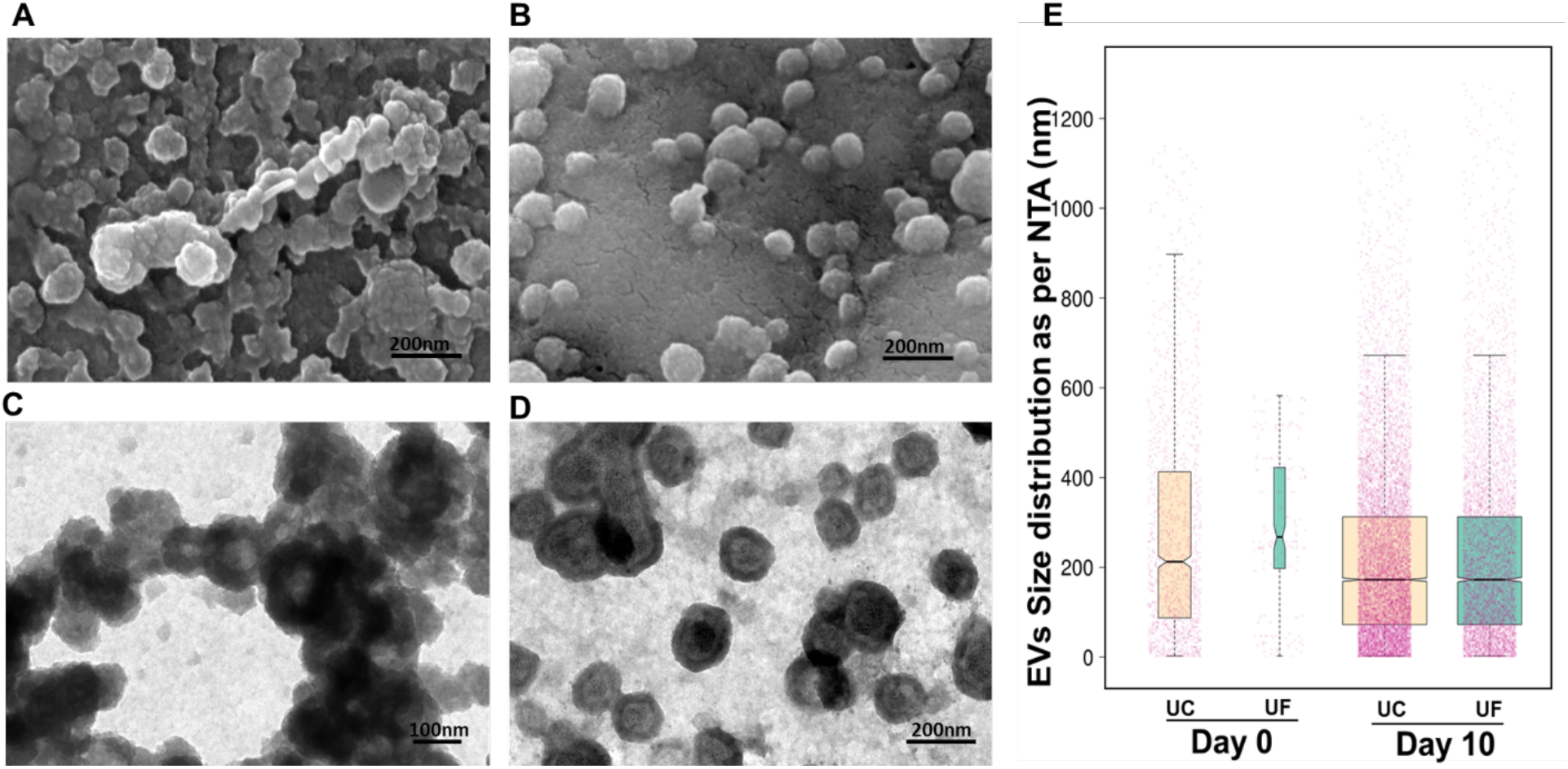
Characterization of blastema associated Exosome like EVs. A and B- FESEM analysis showing surface topology of blastema associated exosome like EVs isolated using ultracentrifugation and ultrafiltration method; C and D- TEM analysis showing structure of blastema associated Exosome like EVs isolated using ultracentrifugation and ultrafiltration method; E- Size distribution of blastema associated Exosome like EVs isolated using ultracentrifugation and ultrafiltration methods.

FESEM analysis revealed that UC isolated EVs appeared as dense, aggregated clusters, whereas UF-isolated EVs were sparsely distributed as individual particles (Figure 2A-B). The dense clustering in UC samples indicated a higher yield compared with UF-isolated samples. However, the vesicles showed irregular surfaces, lacking the smooth and well-defined outline typical for intact exosome like EVs potentially caused by high centrifugal force. In contrast, EVs obtained through UF exhibited smooth and intact surfaces, highlighting better membrane conservation during isolation process. Further TEM analysis supported these findings (Figure 2C-D) by showing similar aggregated clusters in UC isolated EVs, making it challenging to determine single particle structure. Conversely, UF isolated EVs were visible as discrete particles showing typical concave morphology with rim like borders. Although rim-like concave vesicles were observed in UC samples, their clustered arrangement hindered clear individual EV morphology. Moreover, UF isolated particles exhibited central electron dense dot, which may represent protein content which are retained following Folin staining during sample preparation.

Together these observations clarifies that UC yielded a higher quantity of EVs, but the procedure may compromise membrane integrity, whereas UF provides structurally intact vesicles with conserved surface morphology. Thus, the choice of isolation technique should be based on downstream application i.e., UC may be suitable for therapeutic production, while UF is preferable for diagnostic or structural studies where intact morphology is critical.

Moreover, nanoparticle tracking analysis revealed that significant percentage of particles, approximately 85%, fell within the size range associated with exosomes in both methods, further confirming the abundance of EV particles within the sample. Comparison of the results revealed distinct differences in size and concentration between two methods. The UF method yielded vesicles with a median size (X50) of 209.1 nm and a concentration of 43 million particles/mL. On the other hand, the UC method resulted in vesicles with a slightly larger median size of 219.6 nm, accompanied by a higher concentration of 73 million particles/mL. This significant disparity in concentration is supported by our electron microscopy findings. In addition, the measured sizes of the particles likely indicated the mean hydrodynamic diameter of exosomes, and their clusters respectively (Figure. 2E). Consistent with earlier reports, exosomes have been described to range in diameter ranging from 30 - 220 nm. Although the morphology and size distribution were consistent with exosome characteristics, the vesicles were referred to them as exosome like extracellular vesicles (exosome like EVs) in accordance with MISEV guidelines, as their release via the endocytic pathway was not confirmed in this study.

### Exosome like EVs were loaded with more than 200 different types of miRNAs

Isolated exosome like EVs were subjected to RNA extraction through spin column centrifugation to identify the role of exosome like EVs or EV mediated molecules in regeneration. Nanodrop analysis was mainly performed to assess the quantity of the extracted nucleic acid. This analysis provided data on nucleic acid concentration and its absorbance at wavelengths of 260nm and 280nm, as well as the corresponding absorbance ratios (260/280 and 260/230) (see Table S1). The extracted RNA samples underwent Next-Generation Sequencing (NGS) following a quality control (QC) test conducted prior to library preparation and sequencing. The QC report provided information on RNA concentration and RNA Integrity Number (RIN) through an electro phenogram summaries. The outcomes indicated that the total RNA concentration primarily comprises 58% small RNAs, particularly miRNAs, with the remaining fraction comprising other small RNAs. Consequently, the obtained RIN values were low, which was typical for EV derived RNAs (Mussack, Wittmann and Pfaffl, 2019) (see Figure S3). Sequencing of those miRNAs identified around 187 differentially expressed miRNAs wherein 33 miRNAs were upregulated and 21 miRNAs were downregulated (see Table S2). Notably, the downregulated miRNAs dre-miR-455-3p and dre-miR-456 were predicted to target sonic hedgehog (Shh) signalling components. Their reduced expression during regeneration suggests a possible release of miRNA mediated repression on Shh signalling pathways in regenerating tissues. A heatmap illustrating the upregulated and downregulated miRNAs detected in the regenerating zebrafish tissues is shown in Figure 3. The identified DE miRNAs in other regenerating tissues are listed in Table.1.

**Table 1.** The list of some important differentially expressed miRs found in the caudal fin blastema derived exosome like EVs, and their concurrence in other regenerating tissues of zebrafish.

| Differentially expressed miRNAs | Our experiment | Literature | Tissues | References |
| --- | --- | --- | --- | --- |
| Dre-miR-200b | Upregulated | Upregulated | Caudal fin (CF) | (Thatcher *et al.*, 2008) |
| Dre-miR-15b | Upregulated | Upregulated | CF, Heart | (Thatcher *et al.*, 2008; Yin *et al.*, 2008, 2012; King and Yin, 2016) |
| Dre-miR- 21 | Upregulated | Upregulated | CF, Heart, Retina, Nervous tissue | (Thatcher *et al.*, 2008; Yin *et al.*, 2008, 2012; Rajaram *et al.*, 2014; King and Yin, 2016) |
| Dre-let-7e | Upregulated | Downregulated | CF | (Thatcher *et al.*, 2008) |
| Dre-miR-23a | Upregulated | Downregulated | CF, Retina | (Thatcher *et al.*, 2008; Rajaram *et al.*, 2014; King and Yin, 2016) |
| Dre-miR-100 | Upregulated | Downregulated | CF | (Thatcher *et al.*, 2008; Yin *et al.*, 2012) |
| Dre-miR-200c | Downregulated | Downregulated | CF, Heart | (Thatcher *et al.*, 2008; Yin *et al.*, 2012; Beauchemin, Smith and Yin, 2015; King and Yin, 2016) |
| Dre-miR-203b | Upregulated | Downregulated (CF, Retina), Upregulated (Heart) | CF, Heart, Retina | (Rajaram *et al.*, 2014; Beauchemin, Smith and Yin, 2015; King and Yin, 2016) |
| Dre-miR-218a | Upregulated | Upregulated | CF | (King and Yin, 2016) |
| Dre-let-7a | Upregulated | Downregulated | Heart, Retina | (Ramachandran, Fausett and Goldman, 2010; Aguirre *et al.*, 2014; Rajaram *et al.*, 2014) |
| Dre-miR-1 | Upregulated | Downregulated | Heart | (Ceci *et al.*, 2018) |
| Dre-miR-92b | Downregulated | Downregulated | Heart, Nervous Tissue | (Yin *et al.*, 2012; Kim *et al.*, 2018) |
| Dre-miR-146a | Upregulated | Upregulated | Heart, Retina, Nervous Tissue | (Yin *et al.*, 2012; Rajaram *et al.*, 2014; Kim *et al.*, 2018) |
| Dre-miR-146b | Upregulated | Upregulated | Heart, Retina | (Yin *et al.*, 2012; Rajaram *et al.*, 2014) |
| Dre-miR-735 | Upregulated | Upregulated | Heart, Nervous Tissue | (Beauchemin, Smith and Yin, 2015; Kim *et al.*, 2018) |
| Dre-miR-7a | Upregulated | Upregulated | Retina | (Rajaram *et al.*, 2014) |
| Dre-miR-27c | Downregulated | Upregulated | Retina | (Rajaram *et al.*, 2014) |
| Dre-miR-130c | Upregulated | Downregulated | Retina | (Rajaram *et al.*, 2014) |
| Dre-miR-429b | Upregulated | Downregulated | Retina | (Rajaram *et al.*, 2014) |
| Dre-miR-489 | Downregulated | Downregulated | Retina | (Rajaram *et al.*, 2014) |
| Dre-miR-722 | Downregulated | Downregulated | Retina | (Rajaram *et al.*, 2014) |
| Dre-miR-143 | Upregulated | Upregulated | Retina | (Rajaram *et al.*, 2014; Sharma *et al.*, 2019) |
| Dre-miR-142b | Upregulated | Upregulated | Retina | (Rajaram *et al.*, 2014) |
| Dre-miR-738 | Upregulated | Upregulated | Nervous Tissue | (Kim *et al.*, 2018) |
| Dre-miR-196d | Downregulated | stable | Nervous Tissue | (Kim *et al.*, 2018) |

**Figure 3.**
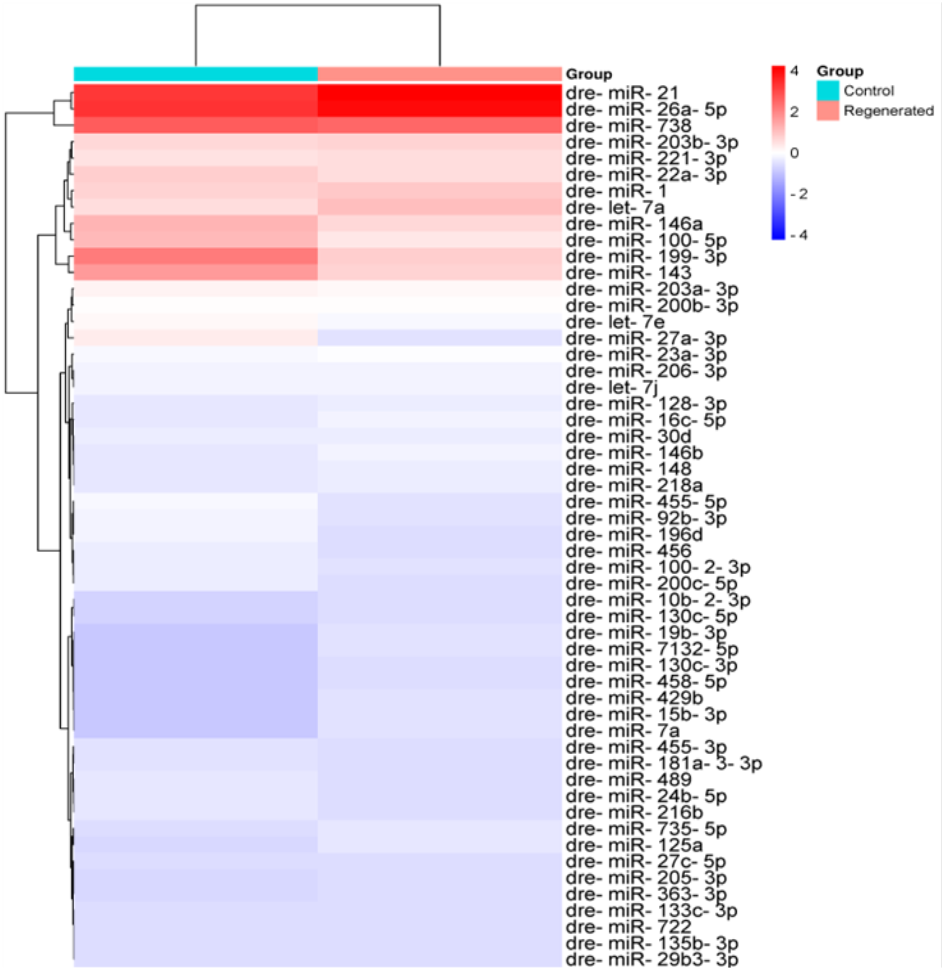
Heatmap analysis of differentially expressed miRNAs extracted from blastema associated exosome like EVs.

## Discussion

Epimorphic regeneration in zebrafish provides an established model to examine the cellular and molecular mechanisms underlying tissue regeneration. While intracellular signalling pathways regulating blastema formation and regeneration in these vertebrates have been studied, mechanisms and factors of coordinated regeneration remain poorly understood. In this context, the present study focuses on exosome like EVs, which are abundantly enriched in regenerating zebrafish caudal fins, particularly within the blastema region.

The regeneration of zebrafish fin progresses through distinct stages, with approximately 10 days post-amputation, the regenerating tissue shows around 30-40% of regeneration which is marked by active blastemal proliferation, differentiation, and tissue patterning (Pfefferli and JaŹwińska, 2015). This phase therefore represents an ideal window to examining molecular signals that coordinate regeneration. In our study, in situ imaging revealed a significant enrichment of exosome like EVs in regenerating fins compared with non-amputated controls, indicating that EV biogenesis is dynamically regulated in response to injury and further suggest that EV accumulation is associated with blastema activity rather than simple tissue remodelling. This observation extends earlier work by Ohgo et al. (2020), who demonstrates EV presence in the regenerating zebrafish fins using in vivo labelling of exosomal markers (Ohgo *et al*., 2020).

Importantly, EV-rich islands were frequently associated with blastema and newly deposited extracellular matrix, including collagen fibre–like structures, suggesting a functional relationship between EV accumulation and tissue morphogenesis. Moreover, spatial analysis revealed preferential enrichment of exosome like EVs around fin rays and joints, with comparatively fewer vesicles in inter-ray membrane regions. This non-uniform distribution suggests that EV-mediated communication is particularly active in regenerative zones associated with bone and supports the indication that epimorphic regeneration involves deep tissue coordination rather than superficial repair processes.

An interesting and potentially novel finding of this study is the organization of EV-associated puncta along filopodia-like membrane extensions within regenerating tissues. Specialized membrane protrusions such as cytonemes and tunnelling nanotubes are well established mediators of long-range intercellular communication and have been implicated in the transport of morphogens and vesicular cargo during embryonic development. In particular, developmental studies have shown that Shh signalling are distributed through filopodia -mediated transport, thereby coordinating tissue patterning across spatially separated cell populations.

In the present study, EV-associated puncta were organized in linear and densely packed arrays along filopodia-like structures, suggesting the possibility of directed EV transport between distal blastemal and proximal regenerating regions. Although direct evidence of EV transport through filopodia in regenerating vertebrate tissues remains limited, these observations suggests that regeneration may utilize communication strategies similar to those employed during development (Kelleher, Fennelly and Rafferty, 2006). Importantly, the identification of EV-associated miRNAs that target developmental signalling pathways including Shh, BMP, and Wnt also strengthens this interpretation. For example, the downregulation of dre-miR-455-3p and dre-miR-456 may accelerate increased Shh expression during regeneration by relieving post-transcriptional repression, thereby supporting blastemal proliferation and tissue patterning. Together, these findings suggest that exosome like EVs may be a regulatory cargo delivered through filopodia-like extensions to coordinate tissue patterning, proliferation, and differentiation during regeneration. This observation is consistent with broader concept that epimorphic regeneration represents, at least in part, a reactivation of developmental programmes. While the present findings are primarily on spatial association and miRNA profiling, future studies involving live imaging, EV tracking, and co-localization analyses of developmental morphogens like Shh will be necessary to establish the mechanistic role of filopodia-mediated EV transport during regeneration.

Furthermore, to explore the molecular composition and potential regulatory functions of exosome like EVs, we examined their nucleic acid cargo with a focus on EV-associated RNAs. Pre sequencing analysis of exosome like EVs cargo revealed a wavy pattern of electrophotography and low RNA integrity number (RIN), consistent with previous reports showing that exosome like EVs are enriched in small regulatory RNAs rather than intact ribosomal RNA (Banigan *et al*., 2013; Barutta *et al*., 2013). Although high RIN values above 8 are generally considered optimal for total cellular RNA, low RIN values are expected and acceptable for EV-derived miRNA preparations. Subsequent miRNA profiling revealed that a regeneration-specific EV miRNA signature was enriched for regulators of cell survival, proliferation, differentiation, inflammation, and tissue patterning, suggesting exosome like EVs as carriers of post-transcriptional regulatory information in regenerating tissues.

Among these, dre-miR-21 shows upregulation in tissues such as the caudal fin, heart, nerve, and retina, playing a role in both neoplastic and non-neoplastic conditions. When dre-miR-21 is downregulated, it leads to increased cell death. While the precise targets of this miRNA remain unidentified, potential candidates include HIF-1α, PTEN, and PDCD4 (Jenike and Halushka, 2021). In neoplastic diseases, miR-21 promotes cell migration through TPM1 and PDCD4, and its upregulation in response to cytokines suggests involvement in inflammation (Liu *et al*., 2014). Moreover, its role in tissue repair have been studied across multiple systems, including wound healing mediated by mesenchymal stem cell (MSC)-derived EVs (Bray *et al*., 2021). Hence, the enrichment of miR-21 within Exosome like EVs suggests a paracrine mechanism by which survival-promoting signals may be distributed from the blastema to surrounding tissues.

Furthermore, few subsets of miRNAs, including dre-miR-200b and dre-miR-100 were prominently expressed in regenerating exosome like EVs. dre-miR-200b hinders tumour growth by disrupting ERK1/2 and AKT signalling pathways, targeting p70S6K1 to suppress HIF-1α expression. miR-200b also increases the sensitivity of H1299 cells to cisplatin while restraining their proliferation and invasion abilities by inhibiting p70S6K1 activity (Jin *et al*., 2020). Overexpression of miR-100 inhibits osteogenic differentiation, while its downregulation enhances the process by targeting BMPR2 (Zeng *et al*., 2012, p. 100). Additionally, miR-100 suppresses the migration and invasion of breast cancer cells by targeting FZD-8 within the canonical Wnt pathway (Jiang *et al*., 2016). Their presence in exosome like EVs supports the hypothesis that regulatory signals that coordinate proliferation and differentiation across spatially distinct regions of regenerating tissue are distributed via vesicle mediated communication.

Additionally, the downregulation of dre-mir-200c targets ZEB1 and ZEB2, both of which suppress E-cadherin expression. Consequently, this downregulation indirectly promotes stronger cell adhesion and intercellular communication. The presence of this miRNA inhibits the transition of cells through epithelial-mesenchymal transition (EMT) and the metastasis of cancer cells. Additionally, it enhances resistance to Anoikis, a form of cell death that occurs when cells lose their attachment to the extracellular matrix (Howe, Cochrane and Richer, 2011). Together, this dynamic interplay of upregulated and downregulated EV-miRNAs likely fine-tunes cellular proliferation, differentiation, and extracellular matrix remodelling.

Recent studies in other regenerative organisms supports functional role of EV mediated signalling in tissue repair. Exosome like EVs derived from regenerating planarian tissues enhance stem cell proliferation and regeneration-associated transcriptional expression, showing the active role of Exosome like EVs in coordinating regeneration in a non-vertebrate model (Avalos Najera, Wong and Forsthoefel, 2025). Similarly, Exosome like EVs derived from urodele amphibians including newts have been shown to promote neurite outgrowth in mammalian neurons, indicating that regenerative EV signals may be evolutionarily conserved (Middleton *et al*., 2023). Although miRNA regulation in axolotl regeneration is studied extensively, direct characterization of axolotl tissue-derived Exosome like EVs during regeneration remains lacking, highlighting an important direction for future investigations (Holman *et al*., 2012).

In mammalian systems, stem, and progenitor cell–derived exosome like EVs have been widely studied in tissue repair and homeostasis. MSC-derived EVs enriched with regulatory miRNAs have been shown to enhance wound healing, angiogenesis, and functional recovery in models of skin, muscle, and cardiac injury (Bhaskara, Anjorin and Wang, 2023; Zhang *et al*., 2023).Together, these studies strengthen the concept that EV-mediated miRNA transfer is a conserved mechanism that promotes tissue repair across taxa and injury contexts.

A major challenge in studying tissue-derived exosome like EVs lies in their isolation from solid tissues. Mechanical or enzymatic dissociation can introduce contaminants and affect vesicle integrity. In the present study, we compared centrifugal filter-based ultrafiltration with conventional ultracentrifugation for isolating exosome like EVs from regenerating zebrafish fins. The observed differences in particle recovery and vesicle integrity underscore the need for optimized and context-specific isolation strategies when studying tissue-derived exosome like EVs.

In summary, this study provides compelling evidence that exosome like EVs are dynamically enriched during zebrafish fin regeneration and carry a regeneration-associated miRNA cargo. our findings support the idea that regeneration involves EV-mediated intercellular communication within regenerating tissues rather than mere intracellular signalling. Collectively, these findings suggest that regenerative tissues may partially recapitulate developmental communication mechanisms through EV-mediated signalling associated with filopodia-like extensions. While the present study establishes a foundational profile of regeneration associated EV-miRNAs, future functional validation studies employing antagomirs, miRNA mimics, or EV uptake inhibition will be essential to define their functional roles. Collectively, these findings highlight the potential importance of regenerating tissue–derived Exosome like EVs as regulators of tissue repair and as promising candidates for regenerative medicine applications.

## Conclusion

This study reports the first evidence on demonstrating the blastema associated exosome like EVs and their potential role in coordinating zebrafish caudal fin regeneration. The abundance of tetraspanin-positive particles is higher in regenerating tissues compared to freshly amputated fins, emphasising the active involvement of EVs in the regenerative process. The isolated tetraspanin-positive particles exhibited characteristic features such as size of about 200 nm and a concave morphology were consistent with exosome-like vesicles. However, since the exosome release via the endocytic pathway was not confirmed in this study, these vesicles are collectively referred to as exosome like EVs or small exosome like EVs according to MISEV guidelines.

Comparative analysis of isolation techniques revealed ultracentrifugation produced a higher yield, it also caused partial membrane disruption, whereas ultrafiltration preserved vesicular morphology and integrity. These observations suggest that the preference of isolation method should be based on the downstream application requirements, for example, ultrafiltration being preferable for therapeutic and structural studies while ultracentrifugation for diagnostic or large-scale preparative purposes.

According to literature, describing the central role of the blastema in regeneration, our findings propose a novel hypothesis that blastema cells may communicate with surrounding and downstream tissues through EV-mediated signalling. These blastema associated exosome like EVs are likely carrying molecular cargo, comprising miRNAs, that control the gene regulatory networks influencing epimorphic regeneration which is limited in mammals. Further marker-based validation and molecular profiling of these vesicles will help reveal their defined role in regeneration signalling.

Altogether, this study highlights blastema-derived exosome like EVs as key mediators of intercellular communication during tissue regeneration and positions them as potential candidates for therapeutic strategies targeting tissue repair, skeletal injuries, and chronic degenerative disorders.

## Methodology

### Caudal fin amputation and quantification of length

Zebrafish (*Danio rerio*) of wildtype were reared and maintained in room temperature for the whole study (Lawrence, 2007). During the experiment, fish were sedated using cold shock method and their pre and post caudal fin amputation length was measured. Amputated fish after recovering from anaesthesia were allowed to regenerate fins (Saxena *et al*., 2012). This experiment was repeated on the 10th day to collect the regenerated fin which were subsequently used for further studies. All experimental procedures involving animals were reviewed and approved by the Institutional Animal Ethical Committee (VIT/IAEC/30/Aug25/25) and were carried out in accordance with established ethical guidelines.

### Whole mount staining and confocal microscopy

Freshly excised control (0 dpa) and regenerated (10 dpa) caudal fins were fixed in Dent’s fixative (1:9 DMSO/methanol, vol/vol) for 24 h. Following fixation, samples were cleared and rehydrated through graded methanol series (75% to 25%) and decalcified using Morse’s solution for 2 h at room temperature, followed by PBS washes. Samples were then treated with 1% KOH and 20% Tween-20 for 1 h at room temperature and permeabilized using diluted Triton X-100. Blocking was performed with 0.05% goat serum in 1% BSA (Alajati *et al*., 2008; Zukor, Kent and Odelberg, 2010; Sakata-Haga *et al*., 2018). Samples were incubated with the primary antibody (Anti-TSPAN9), washed with diluted Triton X-100, and incubated with the secondary antibody (anti-rabbit goat, Abcam), followed by thorough washing. Nuclei were counterstained using DAPI. Finally, fins were mounted in graded glycerol/methanol series (25%, 50%, 75%) and placed on thin coverslips for imaging using a laser scanning confocal microscope (Olympus Fluoview 1000).

### Isolation of exosome like EVs

Exosome like EVs were isolated from amputated zebrafish caudal fins (0dpa and 10dpa) using both ultrafiltration and ultracentrifugation approaches. Initially, fins were homogenized in 2 mM EDTA with PBS, followed by sequential centrifugation at 300 × g for 20 min and 5,000 × g for 20 min at 4°C to remove debris (Vella *et al*., 2017). The supernatant was filtered through a 0.22 μm syringe filter. For ultrafiltration, the filtered solution was concentrated using a Centricon 10 kDa filter tube via centrifugation at 10000 × g for 1 hour at 4°C, with the resulting exosome like EVs stored at -20°C (Sorenson, 2014). For ultracentrifugation, the filtered solution was ultracentrifuged at 100,000 × g for 70 min at 4°C, washed with PBS, and subjected to a second ultracentrifugation at 100,000 × g for 1 hour to concentrate exosome like EVs. The final pellet was resuspended in PBS and stored at -80°C for long-term use.

### Electron microscopical analysis

For microscopical analysis, TEM grids were incubated with 10–20 µL of the EV solution for 1–5 minutes and fixed in 2.5% glutaraldehyde prepared in 0.1 M sodium cacodylate buffer (pH 7.4). The grids were washed three times with cacodylate buffer and stained with an admixture of phosphomolybdic acid and phosphotungstic acid (negative stain) for 2 minutes followed by treatment of UranyLess (positive stain) for 2 minutes. Dehydration was performed using graded ethanol treatments, immersing the grids in 30%, 50%, 70%, 90%, and 100% ethanol for 2 minutes each. The samples were then treated with hexamethyldisilane (HDMS) and dried in a vacuum desiccator. Then, the morphology and surface characteristics of the exosome like EVs were examined using FESEM and TEM.

### Nanoparticle Tracking Analysis (NTA)

Size distribution and concentration of EV samples isolated by ultracentrifugation and ultrafiltration were determined using the ZetaView PMX220 (Particle Metrix) equipped with 550 nm and 580 nm lasers and analysed with Zeta View 8.05.11 SP4 software. EV samples were diluted in PBS to achieve ∼350 particles per frame. Measurements were performed in scatter mode at 29°C across 11 positions in two cycles with the following settings: Sensitivity 80, Shutter 100, Minimal Brightness 30, Trace Length 15, Min Area 10, Max Area 1000 nm/Class 5, Classes per Decade 64, and medium resolution. Camera sensitivity was adjusted to detect dim particles with minimal background noise. Identical settings were maintained for all samples to ensure consistency, and a minimum of three measurements per sample were conducted. Recorded data were processed according to standard settings, and results were interpreted from the generated PDF reports extraction and quantification.

### EV-RNA Extraction, Library Preparation, and Sequencing

Total RNA was extracted from exosome like EVs isolated from control and regenerating zebrafish caudal fins using the Exosomal RNA Isolation Kit (Mini, Norgen Biotek Corp) according to the manufacturer’s spin column protocol. RNA quantity and purity were measured using a Nanodrop Spectrophotometer (Thermo Fisher Scientific), and integrity was evaluated on the Agilent Bioanalyzer 2200.

Libraries for small RNA sequencing were generated from 100 ng of total RNA using the NEXTflex™ Small RNA Sample Preparation protocol (Bio Scientific Corporation, USA). Briefly, RNA molecules were ligated with 3’ and 5’ adapters, reverse transcribed to cDNA, and amplified by 20 PCR cycles of PCR. Amplified libraries were size-selected by polyacrylamide gel electrophoresis, quantified with a Qubit Fluorometer (Thermo Fisher Scientific), and fragment size distribution was confirmed on the Agilent 2100 Bioanalyzer.

Sequencing was performed on the Illumina platform generating 75 bp reads in FASTQ format. Raw reads were processed using srna-workbench V3.0_ALPHA1, including 3’ adapter trimming, length filtering (16–40 bp), and quality filtering (removal of reads with <q30, missing adapters, no insert, or matching other ncRNAs such as rRNA, tRNA, snRNA, snoRNA). An additional 4 bases were trimmed from read starts due to adapter design. Approximately 1.2 million high-quality, non-redundant reads were retained. Reads were aligned using Bowtie 1.1.12 to check for contamination, with unaligned reads used for known and novel miRNA prediction. Conserved miRNAs were identified via homology with zebrafish miRNAs in miRbase, while novel miRNAs were predicted based on secondary structure analysis against the reference genome.

### Differential gene expression (DGE) analysis

Read counts for each miRNA were obtained by taking the number of reads aligning to the respective miRNA sequences. Differential expression analysis was performed using the DESeq package. Library normalization was applied to account for variations in sequencing depth, wherein DESeq calculates a size factor for each sample, and individual read counts were normalized by dividing by this factor. Mean normalized read counts across replicates within a given condition were used for downstream DGE analysis and visualization, including heatmaps generated using ClustVis. For identifying differentially expressed miRNAs, a log2 fold change threshold of ±1 was applied: miRNAs with log2 fold change >1 was considered upregulated, while those with log2 fold change <-1 were considered downregulated.

### Statistical analysis

Quantitative image analysis was performed using ImageJ (National Institutes of Health, Bethesda, MD, USA). Statistical analyses and graphical representations were generated using Shiny Graph (http://shiny.chemgrid.org/boxplotr/).Statistical tests were selected according to the experimental design and distribution of the data. A *P* value of < 0.05 was considered statistically significant.

## Acknowledgement

We acknowledge VIT management for providing TRAship and UGC for providing Savitribai Jyotirao Phule Fellowship to the PhD students involved in this research project. We are also grateful to DST-FIST grant for sponsoring the SEM facility in VIT. Our sincere thanks to Dr. Arunkumar P. from centre for biomaterials cells and molecular theranostics (CBCMT), VIT-Vellore, for letting us use the NTA for our sample analyses and stem cell research centre (SCRC), Christian medical college (CMC), Vellore for allowing us to use their confocal microscopy facility.

