## Supplementary material for "Evidence of Filopodial translocation of Blastema associated microRNA rich Exosome like Extracellular Vesicles": Figure S1, Figure S2, Table S1

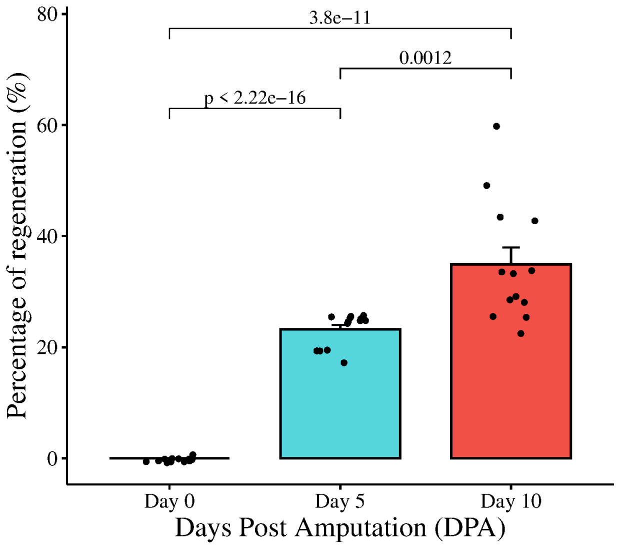

Figure S1. Percentage of caudal fin regeneration in zebrafish

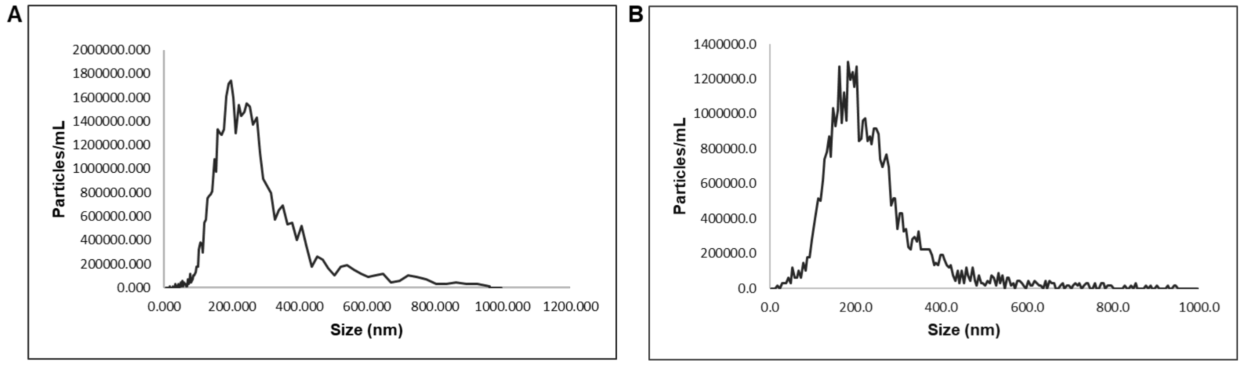

Figure S2. NTA analysis of blastema associated EVs. A and B- Size and concentration of blastema associated EV isolated using ultracentrifugation and ultrafiltration methods.

Table S1. Nanodrop analysis

| Sample name | ng/microliter | A260/A280 | A260/A230 | A260 | A280 | RNA concentration (µg/ml) |
|---|---|---|---|---|---|---|
| Matured fin (Control)) | 253.8 | 2.07 | -2.44 | 6.34 | 3.06 | 12680 |
| 0th Day | 83 | 3.14 | -2.96 | 2.08 | 0.66 | 4160 |
| 10th day (Regenerating fin) | 136.8 | 3.72 | -35.74 | 3.42 | 0.92 | 6840 |

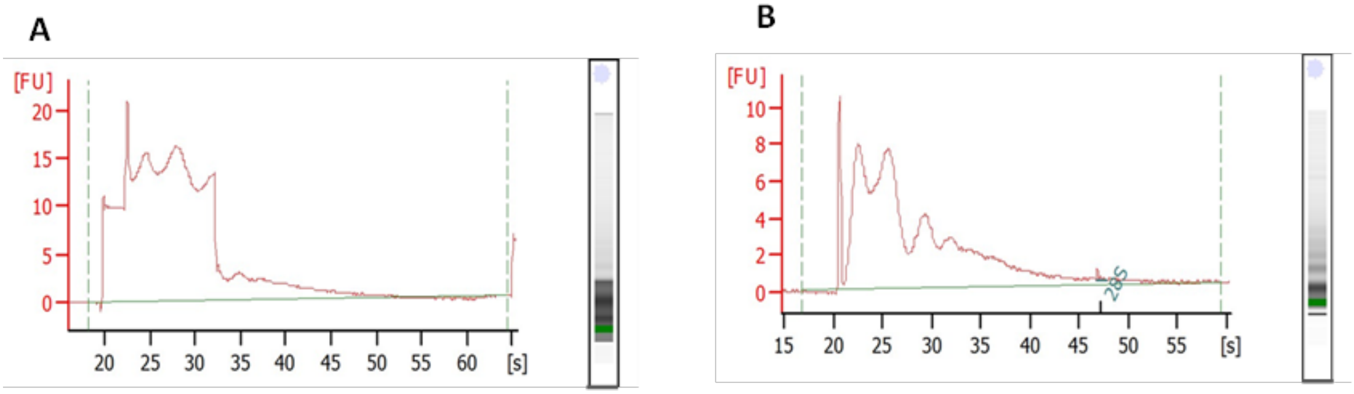


Figure: S3 Electropherogram analysis of EV associated total RNA.

Table: S2 Differentially expressed miRNAs in EV during regeneration.

| miRNA | Day0 | Day10 | Foldchange | Log2Foldchange | Regulation | Accession |
|---|---|---|---|---|---|---|
| dre-miR-21 | 88 | 519 | 5.897727273 | 2.56015911 | UP | MIMAT0001787 Danio rerio miR-21 |
| dre-miR-26a-5p | 90 | 493 | 5.477777778 | 2.45359074 | UP | MIMAT0001794 Danio rerio miR-26a-5p |
| dre-let-7a | 32 | 174 | 5.4375 | 2.442943496 | UP | MIMAT0001759 Danio rerio let-7a |
| dre-miR-738 | 76 | 329 | 4.328947368 | 2.11401626 | UP | MIMAT0003769 Danio rerio miR-738 |
| dre-miR-1 | 36 | 161 | 4.472222222 | 2.160991877 | UP | MIMAT0001768 Danio rerio miR-1 |
| dre-miR-203b-3p | 34 | 135 | 3.970588235 | 1.989352756 | UP | MIMAT0001865 Danio rerio miR-203b-3p |
| dre-miR-221-3p | 30 | 123 | 4.1 | 2.03562391 | UP | MIMAT0001288 Danio rerio miR-221-3p |
| dre-miR-22a-3p | 38 | 120 | 3.157894737 | 1.658963082 | UP | MIMAT0001788 Danio rerio miR-22a-3p |
| dre-miR-429b | 1 | 8 | 8 | 3 | UP | MIMAT0011291 Danio rerio miR-429b |
| dre-miR-200b-3p | 21 | 65 | 3.095238095 | 1.63005039 | UP | MIMAT0001862 Danio rerio miR-200b-3p |
| dre-miR-146a | 47 | 131 | 2.787234043 | 1.47883415 | UP | MIMAT0001843 Danio rerio miR-146a |
| dre-miR-15b-3p | 1 | 7 | 7 | 2.807354922 | UP | MIMAT0031936 Danio rerio miR-15b-3p |
| dre-miR-203a-3p | 24 | 71 | 2.958333333 | 1.564784619 | UP | MIMAT0001278 Danio rerio miR-203a-3p |

| | | | | | | |
|---|---|---|---|---|---|---|
| dre-miR-7a | 1 | 7 | 7 | 2.807354922 | UP | MIMAT0001266 Danio rerio miR-7a |
| dre-miR-199-3p | 65 | 149 | 2.292307692 | 1.196800707 | UP | MIMAT0003155 Danio rerio miR-199-3p |
| dre-miR-143 | 55 | 134 | 2.436363636 | 1.284729477 | UP | MIMAT0001840 Danio rerio miR-143 |
| dre-miR-23a-3p | 18 | 52 | 2.888888889 | 1.530514717 | UP | MIMAT0001790 Danio rerio miR-23a-3p |
| dre-miR-16c-5p | 13 | 39 | 3 | 1.584962501 | UP | MIMAT0001776 Danio rerio miR-16c-5p |
| dre-miR-206-3p | 16 | 41 | 2.5625 | 1.357552005 | UP | MIMAT0001866 Danio rerio miR-206-3p |
| dre-miR-19b-3p | 2 | 8 | 4 | 2 | UP | MIMAT0001783 Danio rerio miR-19b-3p |
| dre-miR-100-5p | 45 | 102 | 2.266666667 | 1.180572246 | UP | MIMAT0001813 Danio rerio miR-100-5p |
| dre-miR-125a | 7 | 21 | 3 | 1.584962501 | UP | MIMAT0001820 Danio rerio miR-125a |
| dre-miR-130c-3p | 1 | 4 | 4 | 2 | UP | MIMAT0001828 Danio rerio miR-130c-3p |
| dre-miR-146b | 13 | 33 | 2.538461538 | 1.343954401 | UP | MIMAT0001844 Danio rerio miR-146b |
| dre-miR-7132-5p | 2 | 7 | 3.5 | 1.807354922 | UP | MIMAT0048669 Danio rerio miR-7132-5p |
| dre-miR-148 | 13 | 32 | 2.461538462 | 1.299560282 | UP | MIMAT0001845 Danio rerio miR-148 |
| dre-miR-218a | 13 | 32 | 2.461538462 | 1.299560282 | UP | MIMAT0001868 Danio rerio miR-218a |
| dre-let-7e | 23 | 47 | 2.043478261 | 1.031026896 | UP | MIMAT0001763 Danio rerio let-7e |
| dre-let-7j | 16 | 36 | 2.25 | 1.169925001 | UP | MIMAT0003015 Danio rerio let-7j |
| dre-miR-458-5p | 1 | 3 | 3 | 1.584962501 | UP | MIMAT0032005 Danio rerio miR-458-5p |
| dre-miR-735-5p | 8 | 19 | 2.375 | 1.247927513 | UP | MIMAT0032012 Danio rerio miR-735-5p |
| dre-miR-30d | 14 | 30 | 2.142857143 | 1.099535674 | UP | MIMAT0001806 Danio rerio miR-30d |
| dre-miR-128-3p | 12 | 25 | 2.083333333 | 1.058893689 | UP | MIMAT0001824 Danio rerio miR-128-3p |

| | | | | | | |
|---|---|---|---|---|---|---|
| dre-let-7b | 16 | 14 | 0.875 | -0.192645078 | NEUTRAL | MIMAT0001760 Danio rerio let-7b |
| dre-let-7c-5p | 14 | 12 | 0.857142857 | -0.222392421 | NEUTRAL | MIMAT0001761 Danio rerio let-7c-5p |
| dre-let-7d-5p | 6 | 10 | 1.666666667 | 0.736965594 | NEUTRAL | MIMAT0001762 Danio rerio let-7d-5p |
| dre-let-7f | 16 | 28 | 1.75 | 0.807354922 | NEUTRAL | MIMAT0001764 Danio rerio let-7f |
| dre-let-7g | 16 | 29 | 1.8125 | 0.857980995 | NEUTRAL | MIMAT0001765 Danio rerio let-7g |
| dre-let-7h | 17 | 17 | 1 | 0 | NEUTRAL | MIMAT0001766 Danio rerio let-7h |
| dre-let-7i | 13 | 8 | 0.615384615 | -0.700439718 | NEUTRAL | MIMAT0001767 Danio rerio let-7i |
| dre-miR-101a | 25 | 33 | 1.32 | 0.40053793 | NEUTRAL | MIMAT0001814 Danio rerio miR-101a |
| dre-miR-103 | 21 | 11 | 0.523809524 | -0.932885804 | NEUTRAL | MIMAT0001816 Danio rerio miR-103 |
| dre-miR-107a-3p | 8 | 8 | 1 | 0 | NEUTRAL | MIMAT0001817 Danio rerio miR-107a-3p |
| dre-miR-10b-5p | 18 | 26 | 1.444444444 | 0.530514717 | NEUTRAL | MIMAT0001268 Danio rerio miR-10b-5p |
| dre-miR-10c-5p | 15 | 8 | 0.533333333 | -0.906890596 | NEUTRAL | MIMAT0001770 Danio rerio miR-10c-5p |
| dre-miR-10d-5p | 11 | 11 | 1 | 0 | NEUTRAL | MIMAT0001771 Danio rerio miR-10d-5p |
| dre-miR-125b-5p | 13 | 16 | 1.230769231 | 0.299560282 | NEUTRAL | MIMAT0001821 Danio rerio miR-125b-5p |
| dre-miR-125c-5p | 9 | 7 | 0.777777778 | -0.362570079 | NEUTRAL | MIMAT0001822 Danio rerio miR-125c-5p |
| dre-miR-126a-3p | 23 | 40 | 1.739130435 | 0.798366139 | NEUTRAL | MIMAT0001823 Danio rerio miR-126a-3p |
| dre-miR-126a-5p | 10 | 14 | 1.4 | 0.485426827 | NEUTRAL | MIMAT0003157 Danio rerio miR-126a-5p |
| dre-miR-126b-3p | 13 | 13 | 1 | 0 | NEUTRAL | MIMAT0011306 Danio rerio miR-126b-3p |
| dre-miR-133a-3p | 30 | 44 | 1.466666667 | 0.552541023 | NEUTRAL | MIMAT0001830 Danio rerio miR-133a-3p |
| dre-miR-133b-3p | 5 | 8 | 1.6 | 0.678071905 | NEUTRAL | MIMAT0001831 Danio rerio miR-133b-3p |
| dre-miR-137-3p | 10 | 5 | 0.5 | -1 | NEUTRAL | MIMAT0001834 Danio rerio miR-137-3p |

| | | | | | | |
|---|---|---|---|---|---|---|
| dre-miR-1388-3p | 13 | 10 | 0.769230769 | -0.378511623 | NEUTRAL | MIMAT0011293 Danio rerio miR-1388-3p |
| dre-miR-1388-5p | 7 | 10 | 1.428571429 | 0.514573173 | NEUTRAL | MIMAT0011292 Danio rerio miR-1388-5p |
| dre-miR-139-5p | 13 | 13 | 1 | 0 | NEUTRAL | MIMAT0003349 Danio rerio miR-139-5p |
| dre-miR-140-3p | 28 | 44 | 1.571428571 | 0.652076697 | NEUTRAL | MIMAT0003159 Danio rerio miR-140-3p |
| dre-miR-140-5p | 14 | 8 | 0.571428571 | -0.807354922 | NEUTRAL | MIMAT0001836 Danio rerio miR-140-5p |
| dre-miR-141-3p | 50 | 50 | 1 | 0 | NEUTRAL | MIMAT0001837 Danio rerio miR-141-3p |
| dre-miR-142a-3p | 5 | 3 | 0.6 | -0.736965594 | NEUTRAL | MIMAT0003160 Danio rerio miR-142a-3p |
| dre-miR-142a-5p | 5 | 5 | 1 | 0 | NEUTRAL | MIMAT0001838 Danio rerio miR-142a-5p |
| dre-miR-142b-5p | 10 | 5 | 0.5 | -1 | NEUTRAL | MIMAT0001839 Danio rerio miR-142b-5p |
| dre-miR-144-5p | 10 | 9 | 0.9 | -0.152003093 | NEUTRAL | MIMAT0031975 Danio rerio miR-144-5p |
| dre-miR-145-3p | 11 | 17 | 1.545454545 | 0.628031223 | NEUTRAL | MIMAT0031976 Danio rerio miR-145-3p |
| dre-miR-145-5p | 12 | 13 | 1.083333333 | 0.115477217 | NEUTRAL | MIMAT0001842 Danio rerio miR-145-5p |
| dre-miR-150 | 14 | 9 | 0.642857143 | -0.637429921 | NEUTRAL | MIMAT0001846 Danio rerio miR-150 |
| dre-miR-152 | 18 | 14 | 0.777777778 | -0.362570079 | NEUTRAL | MIMAT0001847 Danio rerio miR-152 |
| dre-miR-153c-3p | 4 | 3 | 0.75 | -0.415037499 | NEUTRAL | MIMAT0001850 Danio rerio miR-153c-3p |
| dre-miR-155 | 13 | 11 | 0.846153846 | -0.2410081 | NEUTRAL | MIMAT0001851 Danio rerio miR-155 |
| dre-miR-15a-5p | 9 | 14 | 1.555555556 | 0.637429921 | NEUTRAL | MIMAT0001772 Danio rerio miR-15a-5p |
| dre-miR-15b-5p | 12 | 6 | 0.5 | -1 | NEUTRAL | MIMAT0001773 Danio rerio miR-15b-5p |
| dre-miR-15c | 9 | 14 | 1.555555556 | 0.637429921 | NEUTRAL | MIMAT0003764 Danio rerio miR-15c |

| | | | | | | |
|---|---|---|---|---|---|---|
| dre-miR-16a | 9 | 7 | 0.777777778 | -0.362570079 | NEUTRAL | MIMAT0001774 Danio rerio miR-16a |
| dre-miR-16b | 15 | 27 | 1.8 | 0.847996907 | NEUTRAL | MIMAT0001775 Danio rerio miR-16b |
| dre-miR-16c-3p | 9 | 8 | 0.888888889 | -0.169925001 | NEUTRAL | MIMAT0031937 Danio rerio miR-16c-3p |
| dre-miR-17a-5p | 13 | 9 | 0.692307692 | -0.530514717 | NEUTRAL | MIMAT0001777 Danio rerio miR-17a-5p |
| dre-miR-181a-2-3p | 8 | 5 | 0.625 | -0.678071905 | NEUTRAL | MIMAT0032007 Danio rerio miR-181a-2-3p |
| dre-miR-181a-3p | 12 | 6 | 0.5 | -1 | NEUTRAL | MIMAT0001282 Danio rerio miR-181a-3p |
| dre-miR-181a-5-3p | 10 | 10 | 1 | 0 | NEUTRAL | MIMAT0048655 Danio rerio miR-181a-5-3p |
| dre-miR-181a-5p | 34 | 37 | 1.088235294 | 0.121990524 | NEUTRAL | MIMAT0001623 Danio rerio miR-181a-5p |
| dre-miR-181b-5p | 13 | 10 | 0.769230769 | -0.378511623 | NEUTRAL | MIMAT0001270 Danio rerio miR-181b-5p |
| dre-miR-181c-5p | 7 | 4 | 0.571428571 | -0.807354922 | NEUTRAL | MIMAT0001852 Danio rerio miR-181c-5p |
| dre-miR-182-5p | 12 | 14 | 1.166666667 | 0.222392421 | NEUTRAL | MIMAT0001271 Danio rerio miR-182-5p |
| dre-miR-183-5p | 15 | 9 | 0.6 | -0.736965594 | NEUTRAL | MIMAT0001273 Danio rerio miR-183-5p |
| dre-miR-184 | 13 | 23 | 1.769230769 | 0.823122238 | NEUTRAL | MIMAT0001853 Danio rerio miR-184 |
| dre-miR-187 | 9 | 7 | 0.777777778 | -0.362570079 | NEUTRAL | MIMAT0001274 Danio rerio miR-187 |
| dre-miR-18c | 8 | 4 | 0.5 | -1 | NEUTRAL | MIMAT0001781 Danio rerio miR-18c |
| dre-miR-192 | 9 | 6 | 0.666666667 | -0.584962501 | NEUTRAL | MIMAT0001275 Danio rerio miR-192 |
| dre-miR-193a-5p | 10 | 5 | 0.5 | -1 | NEUTRAL | MIMAT0031981 Danio rerio miR-193a-5p |
| dre-miR-193b-3p | 12 | 8 | 0.666666667 | -0.584962501 | NEUTRAL | MIMAT0001857 Danio rerio miR-193b-3p |
| dre-miR-194a | 5 | 6 | 1.2 | 0.263034406 | NEUTRAL | MIMAT0001858 Danio rerio miR-194a |

| | | | | | | |
|---|---|---|---|---|---|---|
| dre-miR-196a-3p | 5 | 4 | 0.8 | -0.321928095 | NEUTRAL | MIMAT0031983 Danio rerio miR-196a-3p |
| dre-miR-196a-5p | 16 | 9 | 0.5625 | -0.830074999 | NEUTRAL | MIMAT0001276 Danio rerio miR-196a-5p |
| dre-miR-196b | 7 | 12 | 1.714285714 | 0.777607579 | NEUTRAL | MIMAT0001860 Danio rerio miR-196b |
| dre-miR-199-3-3p | 62 | 86 | 1.387096774 | 0.472068444 | NEUTRAL | MIMAT0031922 Danio rerio miR-199-3-3p |
| dre-miR-199-5p | 20 | 33 | 1.65 | 0.722466024 | NEUTRAL | MIMAT0001277 Danio rerio miR-199-5p |
| dre-miR-19d-3p | 8 | 5 | 0.625 | -0.678071905 | NEUTRAL | MIMAT0001785 Danio rerio miR-19d-3p |
| dre-miR-200a-3p | 25 | 19 | 0.76 | -0.395928676 | NEUTRAL | MIMAT0001861 Danio rerio miR-200a-3p |
| dre-miR-200a-5p | 11 | 8 | 0.727272727 | -0.459431619 | NEUTRAL | MIMAT0031984 Danio rerio miR-200a-5p |
| dre-miR-200b-5p | 9 | 8 | 0.888888889 | -0.169925001 | NEUTRAL | MIMAT0031985 Danio rerio miR-200b-5p |
| dre-miR-200c-3p | 29 | 31 | 1.068965517 | 0.096215315 | NEUTRAL | MIMAT0001863 Danio rerio miR-200c-3p |
| dre-miR-203b-5p | 9 | 11 | 1.222222222 | 0.289506617 | NEUTRAL | MIMAT0003407 Danio rerio miR-203b-5p |
| dre-miR-204-5p | 23 | 19 | 0.826086957 | -0.275634443 | NEUTRAL | MIMAT0001279 Danio rerio miR-204-5p |
| dre-miR-205-5p | 29 | 46 | 1.586206897 | 0.665580961 | NEUTRAL | MIMAT0001280 Danio rerio miR-205-5p |
| dre-miR-20a-5p | 16 | 9 | 0.5625 | -0.830074999 | NEUTRAL | MIMAT0001786 Danio rerio miR-20a-5p |
| dre-miR-210-5p | 12 | 6 | 0.5 | -1 | NEUTRAL | MIMAT0003392 Danio rerio miR-210-5p |
| dre-miR-214 | 35 | 32 | 0.914285714 | -0.129283017 | NEUTRAL | MIMAT0001283 Danio rerio miR-214 |
| dre-miR-2184 | 14 | 10 | 0.714285714 | -0.485426827 | NEUTRAL | MIMAT0011288 Danio rerio miR-2184 |
| dre-miR-221-5p | 14 | 7 | 0.5 | -1 | NEUTRAL | MIMAT0031926 Danio rerio miR-221-5p |
| dre-miR-222a-3p | 25 | 40 | 1.6 | 0.678071905 | NEUTRAL | MIMAT0001289 Danio rerio miR-222a-3p |

| | | | | | | |
|---|---|---|---|---|---|---|
| dre-miR-222a-5p | 17 | 21 | 1.235294118 | 0.304854582 | NEUTRAL | MIMAT0031927 Danio rerio miR-222a-5p |
| dre-miR-222b | 9 | 12 | 1.333333333 | 0.415037499 | NEUTRAL | MIMAT0011303 Danio rerio miR-222b |
| dre-miR-223 | 12 | 16 | 1.333333333 | 0.415037499 | NEUTRAL | MIMAT0001290 Danio rerio miR-223 |
| dre-miR-22a-5p | 15 | 20 | 1.333333333 | 0.415037499 | NEUTRAL | MIMAT0031942 Danio rerio miR-22a-5p |
| dre-miR-22b-3p | 14 | 7 | 0.5 | -1 | NEUTRAL | MIMAT0001789 Danio rerio miR-22b-3p |
| dre-miR-22b-5p | 5 | 3 | 0.6 | -0.736965594 | NEUTRAL | MIMAT0031943 Danio rerio miR-22b-5p |
| dre-miR-23a-3-5p | 8 | 7 | 0.875 | -0.192645078 | NEUTRAL | MIMAT0031945 Danio rerio miR-23a-3-5p |
| dre-miR-23b-3p | 10 | 16 | 1.6 | 0.678071905 | NEUTRAL | MIMAT0048661 Danio rerio miR-23b-3p |
| dre-miR-24 | 17 | 25 | 1.470588235 | 0.556393349 | NEUTRAL | MIMAT0001792 Danio rerio miR-24 |
| dre-miR-24b-3p | 10 | 14 | 1.4 | 0.485426827 | NEUTRAL | MIMAT0048663 Danio rerio miR-24b-3p |
| dre-miR-25-3p | 22 | 34 | 1.545454545 | 0.628031223 | NEUTRAL | MIMAT0001793 Danio rerio miR-25-3p |
| dre-miR-26a-3p | 6 | 5 | 0.833333333 | -0.263034406 | NEUTRAL | MIMAT0031947 Danio rerio miR-26a-3p |
| dre-miR-26b | 122 | 178 | 1.459016393 | 0.544996093 | NEUTRAL | MIMAT0001795 Danio rerio miR-26b |
| dre-miR-27a-5p | 12 | 7 | 0.583333333 | -0.777607579 | NEUTRAL | MIMAT0031949 Danio rerio miR-27a-5p |
| dre-miR-27b-3p | 26 | 39 | 1.5 | 0.584962501 | NEUTRAL | MIMAT0001797 Danio rerio miR-27b-3p |
| dre-miR-27b-5p | 8 | 9 | 1.125 | 0.169925001 | NEUTRAL | MIMAT0031950 Danio rerio miR-27b-5p |
| dre-miR-27c-3p | 29 | 58 | 2 | 1 | NEUTRAL | MIMAT0001798 Danio rerio miR-27c-3p |
| dre-miR-27d | 12 | 8 | 0.666666667 | -0.584962501 | NEUTRAL | MIMAT0001799 Danio rerio miR-27d |
| dre-miR-27e | 18 | 16 | 0.888888889 | -0.169925001 | NEUTRAL | MIMAT0001800 Danio rerio miR-27e |

| | | | | | | |
|---|---|---|---|---|---|---|
| dre-miR-29a | 9 | 8 | 0.888888889 | -0.169925001 | NEUTRAL | MIMAT0001802 Danio rerio miR-29a |
| dre-miR-30b | 11 | 13 | 1.181818182 | 0.2410081 | NEUTRAL | MIMAT0001804 Danio rerio miR-30b |
| dre-miR-30c-5p | 16 | 26 | 1.625 | 0.700439718 | NEUTRAL | MIMAT0001805 Danio rerio miR-30c-5p |
| dre-miR-30e-3p | 16 | 11 | 0.6875 | -0.540568381 | NEUTRAL | MIMAT0003402 Danio rerio miR-30e-3p |
| dre-miR-30e-5p | 12 | 15 | 1.25 | 0.321928095 | NEUTRAL | MIMAT0001807 Danio rerio miR-30e-5p |
| dre-miR-31 | 12 | 10 | 0.833333333 | -0.263034406 | NEUTRAL | MIMAT0003347 Danio rerio miR-31 |
| dre-miR-338-3p | 10 | 6 | 0.6 | -0.736965594 | NEUTRAL | MIMAT0048673 Danio rerio miR-338-3p |
| dre-miR-365 | 12 | 6 | 0.5 | -1 | NEUTRAL | MIMAT0001875 Danio rerio miR-365 |
| dre-miR-375 | 16 | 15 | 0.9375 | -0.093109404 | NEUTRAL | MIMAT0001876 Danio rerio miR-375 |
| dre-miR-429a | 16 | 12 | 0.75 | -0.415037499 | NEUTRAL | MIMAT0001624 Danio rerio miR-429a |
| dre-miR-451 | 16 | 29 | 1.8125 | 0.857980995 | NEUTRAL | MIMAT0001634 Danio rerio miR-451 |
| dre-miR-454a | 9 | 18 | 2 | 1 | NEUTRAL | MIMAT0001877 Danio rerio miR-454a |
| dre-miR-454b | 22 | 18 | 0.818181818 | -0.289506617 | NEUTRAL | MIMAT0001878 Danio rerio miR-454b |
| dre-miR-455-2-5p | 8 | 6 | 0.75 | -0.415037499 | NEUTRAL | MIMAT0011302 Danio rerio miR-455-2-5p |
| dre-miR-457a | 13 | 8 | 0.615384615 | -0.700439718 | NEUTRAL | MIMAT0001883 Danio rerio miR-457a |
| dre-miR-457b-5p | 7 | 10 | 1.428571429 | 0.514573173 | NEUTRAL | MIMAT0001884 Danio rerio miR-457b-5p |
| dre-miR-458-3p | 16 | 12 | 0.75 | -0.415037499 | NEUTRAL | MIMAT0001885 Danio rerio miR-458-3p |
| dre-miR-460-5p | 9 | 7 | 0.777777778 | -0.362570079 | NEUTRAL | MIMAT0001887 Danio rerio miR-460-5p |
| dre-miR-462 | 17 | 13 | 0.764705882 | -0.387023123 | NEUTRAL | MIMAT0001855 Danio rerio miR-462 |

| | | | | | | |
|---|---|---|---|---|---|---|
| dre-miR-499-5p | 10 | 13 | 1.3 | 0.378511623 | NEUTRAL | MIMAT0003749 Danio rerio miR-499-5p |
| dre-miR-7133-3p | 6 | 5 | 0.833333333 | -0.263034406 | NEUTRAL | MIMAT0048665 Danio rerio miR-7133-3p |
| dre-miR-7147 | 10 | 7 | 0.7 | -0.514573173 | NEUTRAL | MIMAT0028205 Danio rerio miR-7147 |
| dre-miR-724 | 4 | 8 | 2 | 1 | NEUTRAL | MIMAT0003752 Danio rerio miR-724 |
| dre-miR-725-3p | 27 | 44 | 1.62962963 | 0.704544116 | NEUTRAL | MIMAT0003753 Danio rerio miR-725-3p |
| dre-miR-725-5p | 7 | 6 | 0.857142857 | -0.222392421 | NEUTRAL | MIMAT0032010 Danio rerio miR-725-5p |
| dre-miR-727-3p | 4 | 4 | 1 | 0 | NEUTRAL | MIMAT0003756 Danio rerio miR-727-3p |
| dre-miR-731 | 11 | 13 | 1.181818182 | 0.2410081 | NEUTRAL | MIMAT0003761 Danio rerio miR-731 |
| dre-miR-733 | 8 | 5 | 0.625 | -0.678071905 | NEUTRAL | MIMAT0003763 Danio rerio miR-733 |
| dre-miR-735-3p | 9 | 8 | 0.888888889 | -0.169925001 | NEUTRAL | MIMAT0003766 Danio rerio miR-735-3p |
| dre-miR-737-5p | 2 | 4 | 2 | 1 | NEUTRAL | MIMAT0032013 Danio rerio miR-737-5p |
| dre-miR-7b | 7 | 10 | 1.428571429 | 0.514573173 | NEUTRAL | MIMAT0001265 Danio rerio miR-7b |
| dre-miR-92a-3p | 55 | 89 | 1.618181818 | 0.694373717 | NEUTRAL | MIMAT0001808 Danio rerio miR-92a-3p |
| dre-miR-93 | 10 | 17 | 1.7 | 0.765534746 | NEUTRAL | MIMAT0001810 Danio rerio miR-93 |
| dre-miR-9-3p | 4 | 5 | 1.25 | 0.321928095 | NEUTRAL | MIMAT0003156 Danio rerio miR-9-3p |
| dre-miR-9-5p | 9 | 7 | 0.777777778 | -0.362570079 | NEUTRAL | MIMAT0001769 Danio rerio miR-9-5p |
| dre-miR-99 | 44 | 69 | 1.568181818 | 0.649092838 | NEUTRAL | MIMAT0001812 Danio rerio miR-99 |
| dre-miR-196d | 16 | 3 | 0.1875 | -2.415037499 | DOWN | MIMAT0011313 Danio rerio miR-196d |
| dre-miR-200c-5p | 15 | 4 | 0.266666667 | -1.906890596 | DOWN | MIMAT0031986 Danio rerio miR-200c-5p |

| | | | | | | |
|---|---|---|---|---|---|---|
| dre-miR-455-5p | 17 | 6 | 0.352941176 | -1.502500341 | DOWN | MIMAT0001879 Danio rerio miR-455-5p |
| dre-miR-489 | 12 | 3 | 0.25 | -2 | DOWN | MIMAT0002940 Danio rerio miR-489 |
| dre-miR-216b | 12 | 4 | 0.333333333 | -1.584962501 | DOWN | MIMAT0001867 Danio rerio miR-216b |
| dre-miR-456 | 14 | 5 | 0.357142857 | -1.485426827 | DOWN | MIMAT0001882 Danio rerio miR-456 |
| dre-miR-135b-3p | 9 | 3 | 0.333333333 | -1.584962501 | DOWN | MIMAT0032006 Danio rerio miR-135b-3p |
| dre-miR-92b-3p | 16 | 7 | 0.4375 | -1.192645078 | DOWN | MIMAT0001809 Danio rerio miR-92b-3p |
| dre-miR-29b3-3p | 9 | 3 | 0.333333333 | -1.584962501 | DOWN | MIMAT0048668 Danio rerio miR-29b3-3p |
| dre-miR-27a-3p | 27 | 13 | 0.481481481 | -1.054447784 | DOWN | MIMAT0001796 Danio rerio miR-27a-3p |
| dre-miR-130c-5p | 5 | 1 | 0.2 | -2.321928095 | DOWN | MIMAT0031969 Danio rerio miR-130c-5p |
| dre-miR-722 | 9 | 3 | 0.333333333 | -1.584962501 | DOWN | MIMAT0003748 Danio rerio miR-722 |
| dre-miR-24b-5p | 12 | 5 | 0.416666667 | -1.263034406 | DOWN | MIMAT0048662 Danio rerio miR-24b-5p |
| dre-miR-181a-3-3p | 10 | 4 | 0.4 | -1.321928095 | DOWN | MIMAT0048654 Danio rerio miR-181a-3-3p |
| dre-miR-100-2-3p | 15 | 7 | 0.466666667 | -1.099535674 | DOWN | MIMAT0031958 Danio rerio miR-100-2-3p |
| dre-miR-27c-5p | 8 | 3 | 0.375 | -1.415037499 | DOWN | MIMAT0003401 Danio rerio miR-27c-5p |
| dre-miR-455-3p | 11 | 5 | 0.454545455 | -1.137503524 | DOWN | MIMAT0031993 Danio rerio miR-455-3p |
| dre-miR-205-3p | 7 | 3 | 0.428571429 | -1.222392421 | DOWN | MIMAT0031925 Danio rerio miR-205-3p |
| dre-miR-10b-2-3p | 5 | 2 | 0.4 | -1.321928095 | DOWN | MIMAT0031934 Danio rerio miR-10b-2-3p |
| dre-miR-133c-3p | 9 | 4 | 0.444444444 | -1.169925001 | DOWN | MIMAT0001832 Danio rerio miR-133c-3p |
| dre-miR-363-3p | 7 | 3 | 0.428571429 | -1.222392421 | DOWN | MIMAT0001874 Danio rerio miR-363-3p |
